# Overcoming acquired doxorubicin resistance of ovarian carcinoma cells by verapamil-mediated promotion of DNA damage-driven cell death

**DOI:** 10.1101/2025.04.23.650268

**Authors:** Elvira Mukinovic, Sina Federmann, Larissa Meßling, Marlena Sekeres, Julia Mann, Lena Abbey, Matthias U. Kassack, Gerhard Fritz

**Author notes:** **Corresponding author:** Gerhard Fritz, PhD, Institute of Toxicology Medical Faculty and University Hospital Heinrich Heine University Duesseldorf 40225 Duesseldorf, Germany.

## Abstract

The therapeutic efficacy of anticancer therapeutics is limited by acquired drug resistance of tumor cells. Here, we aim to characterize and overcome resistance mechanisms of ovarian cancer cells to the anthracycline derivative doxorubicin (Doxo). To this end, comparative analyses of Doxo-induced stress responses of parental A2780 and Doxo-resistant A2780ADR variant were performed. A2780ADR cells revealed cross-resistance to multiple compounds, including anticancer drugs (cisplatin (CisPt), etoposide (Eto)) and DNA repair/ DNA damage response (DDR) inhibitors (olaparib, niraparib, entinostat, prexasertib, rabusertib). A2780ADR cells formed significantly less DNA double-strand breaks (DSB) after Doxo exposure as compared to A2780, resulting in a mitigated DDR, reduced proliferation inhibition and attenuated apoptosis. Potential resistance mechanisms identified to contribute to Doxo resistance of A2780ADR cells include increased Doxo efflux due to increased MDR1 expression and reduced topoisomerase IIα protein expression. Substantial resensitization of A2780ADR cells to Doxo was achieved by both the RAC1 GTPase inhibitor EHT1864, the histone deacetylase inhibitor entinostat (Est) and, most effectively, the calcium channel blocker verapamil (Ver). Notably, Ver-mediated sensitization also pertains to Eto and CisPt. The synergistic effect of Ver in combination with Doxo, which is reflected by low combination index (CI), likely involves inhibition of MDR1-mediated drug transport, leading to increased intracellular steady state levels of Doxo and elevated DNA damage formation, eventually promoting pro-apoptotic DDR. However, combination treatment with Doxo and Ver also increased the cytotoxic response of non-malignant murine cardiomyocytes (HL-1), murine embryonic stem cells (mESC) and human induced pluripotent stem cells (hiPSC). Based on our data we suggest inhibition of MDR1-mediated Doxo efflux by Ver as a useful approach to overcome acquired drug resistance of A2780ADR cells because it promotes DDR-related pro-death mechanisms, yet at the price of a potentially increased risk of normal tissue toxicity.

**Graphical abstract:**
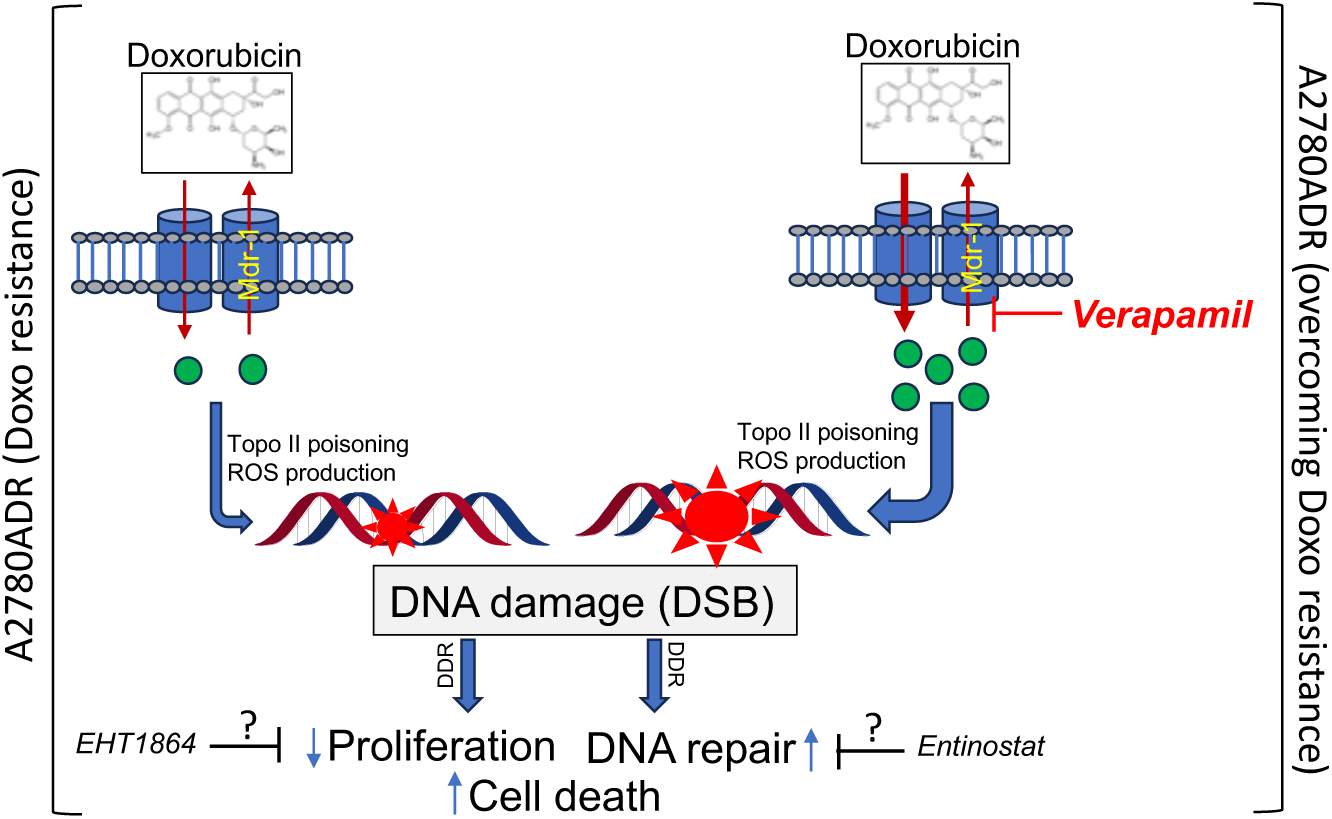
Verapamil-mediated resensitization of Doxo-resistant tumor cells. As concluded from our data we suggest that verapamil increases the anticancer efficacy of doxorubicin (Doxo) in a synergistic manner in anticancer drug resistant ovarian A2780ADR cells. This is likely due to inhibition of MDR1-mediated drug export by Ver, leading to higher intracellular steady-state concentrations of Doxo. In consequence, Doxo-mediated Topo II poisoning is promoted, eventually causing increased DNA damage (DSB) formation and activation of DDR-related signaling mechanims, which in turn impair cell proliferation and stimulate cell death. Apart from verapamil, inhibition of Rac1 GTPase-regulated signaling by EHT1864 and inhibition of HDACs class I by the HDACi entinostat are also useful to overcome Doxo resistance of A2780ADR cells, yet with the exact molecular mechanisms involved being still unclear.

## Introduction

The anticancer efficacy of tumor therapeutics is impaired by inherent or acquired tumor cell resistance. In addition, adverse effects on normal tissue limit the maximum possible cumulative dose of the anticancer drug that can be applied. Against this background, alternative and well-tolerated therapeutic options are needed. Anthracyclines are conventional (i.e. genotoxic) anticancer therapeutics (cAT) which are used for the treatment of numerous malignancies, including hematological disorders, sarcomas and breast cancer [1]. They impair the genetic integrity and thus the malignancy of tumor cells by inhibiting topoisomerase II (Topo II), which is essential for DNA replication. In consequence of Topo II poisoning, DNA double-strand breaks (DSB) are formed, which effectively trigger mechanisms of cell death [2, 3]. DNA intercalation, inhibition of DNA helicases, disruption of mitochondrial functions and formation of reactive oxygen species (ROS) [4] also contribute to the antitumor effect of anthracyclines. Tumor cell resistance mechanisms are often agent-specific and were classified into pre-, on– and post-target mechanisms [5]. Pre-target resistance mechanisms, such as mechanisms of transport or detoxification, eventually reduce the level of drug-induced primary DNA damage and, in consequence, DNA damage-triggered cell death. Regarding Doxo, overexpression of the drug exporter protein p-glycoprotein (P-gp/MDR1) is considered as an important mechanism of acquired Doxo resistance [6, 7]. However, since cells express a variety of different transporters (importers and exporters) for Doxo and other cAT [7, 8], the outcome of anticancer drug treatment is ultimately determined by the combined activity/expression of multiple importers and exporters. Against this background and having in mind that cellular mechanisms contributing to acquired drug resistance in a genetically heterogenous tumor cell population are likely manifold, it would be desirable to effectively target transport dependent (i.e. pre-target mechanisms) and/or transport-independent (i.e. on-or post-target) mechanisms that contribute to drug resistance. Since cAT-induced DNA damage induces a complex stress response termed DNA damage response (DDR), which regulates mechanisms of cell cycle progression, DNA repair and, finally, survival– and death-related pathways [3, 9], factors of the DDR are considered as particular promising pharmacological targets to overcome inherent or acquired tumor cell resistance [10–12].

The DNA damage response (DDR) gets fine-tuned by the PI3-like kinase Ataxia telangiectasia mutated (ATM) and the ATM and Rad3-related kinase (ATR) [13–15], with ATM being of particular relevance for the regulation of DSB-induced stress responses and ATR for replicative stress responses [16–18]. By coordinating the activation of cell cycle checkpoints, DNA repair and cell death-related pathways, the ATM/ATR-regulated network represents the major molecular switch that defines the balance between survival and death [3]. In line with this, ATM– and ATR-regulated pathways contribute to tumor cell resistance [19] and DDR modulating compounds have been proven as useful to improve anticancer therapy [20–23]. In case of oncogene-driven replicative stress, tumor cells are often particular sensitive to compounds that impair a coordinated replicative stress response, thereby eventually enforcing replication fork collapse and death [21, 22, 24–27]. Alterations in DNA repair provides another Achilleś heel for personalized anticancer therapy as reflected by synthetic lethality [28–30]. Here, defective DSB repair by homologous recombination (HR), for example due to hereditary BRCA1/2 deficiency (BRCAness), predicts the malignant cellś hypersensitivity to simultaneous inhibition of PARP-related backup DNA repair pathways by PARP inhibitors (e.g. olaparib or niraparib) [31, 32].

In our study, we used ovarian carcinoma cells (A2780ADR) as *in vitro* model system of acquired Doxo resistance [33]. We comparatively characterized (i) the stress responses of A2780 parental and A2780ADR cells to Doxo treatment, (ii) the cross-resistance of A2780ADR to other anticancer drugs (Eto, CisPt) as well as to a set of candidate compounds interfering with DDR / DNA repair, RAC1 GTPase signaling or drug transport and (iii) the outcome of a combined treatment of A2780ADR cells with Doxo plus the aforementioned inhibitors. Thereby, we aimed to identify compounds that are particularly effective to overcome aquired Doxo resistance.

## Materials and Methods

### Materials

Chemicals were obtained from the following providers: Entinostat (MS-275) was obtained from Selleck Chemicals (Houston, USA), Doxorubicin from STADA Consumer Health & STADAPHARM GmbH (Bad Vilbel, Germany), Etoposide, Ehop16, Prexasertib (AZD-7762) and Dexrazoxan are from Sigma-Aldrich (St. Louis, USA), Cisplatin from Accord Healthcare GmbH (München, Germany), Olaparib from Apexbio Technology LLC (Houston, USA), Niraparib from MedChemExpress LLC (Monmouth Junction, USA), EHT1864 was purchased from Tocris Bioscience (Bristol, United Kingdom), Rabusertib (LY2603618), Ricolinostat (ACY-1215) from MedChemExpress LLC (Monmouth Junction, USA) and Verapamil from Thermo Fisher Scientific (Waltham, USA). The following primary antibodies were used: ATP7A, ERK2, phospho-Histone H3 (Ser10) from Thermo Fisher Scientific Inc. (Carlsbad, USA), Caspase-7 cleaved (Asp198), pChk1 (Ser 345), Cyclin B1, Galactosidase beta (E2U2I), GAPDH (14C10), MDR1 /ABCB1 (D3H1Q), pP53 (S15), PARP, TopBP1(D8G4L), Topoisomerase II alpha (D10G9), 53BP1, Ki67 from Cell Signaling Technology Inc. (Danvers, USA), pChk2 (T68) [Y171], CTR1 / SLC31A1 [EPR7936], Rad51 from Abcam plc. (Bath, UK), γH2AX (Ser 139) clone JBW301, pKAP-1 (S824), pRPA32 (S4/S8) phospho from Bethyl Laboratories Inc. (Montgomery, USA), OCT2 from Biozol Diagnostics Vertrieb GmbH (Eching, Germany), p16 (F-12) and p21 (C-19) from Santa Cruz Biotechnology (Dallas, USA). As secondary antibodies horseradish peroxidase-conjugated secondary antibodies goat anti-mouse IgG and mouse anti-rabbit IgG were used (Rockland, Limerick, PA, USA).

### Cell culture and treatment of cells

Parental A2780 ovarian carcinoma cells (A2780) as well as doxorubicin resistant variant (A2780ADR) originate from the European Collection of Authenticated Cell Cultures (ECACC, Wiltshire, England) and were cultured in RPMI-1640 medium (Sigma-Aldrich, St. Louis, USA) containing 10% fetal calf serum, 1% glutamine and 1% Pen/Strep at 37 °C in a humidified atmosphere containing 5% CO_2_. Cells were authenticated by STR profiling during the last three years. All experiments were performed with mycoplasma free cells. Immortalized HL-1 cardiomyocytes were provided by W.C. Claycomb (New Orleans, LA, USA) [34] and were grown on gelatin (2 mg/ml)/fibronectin (1mg/ml) (Sigma Aldrich, Darmstadt, Germany) coated dishes and maintained in Claycomb medium, supplemented with 10% FBS and 100 μM norepinephrine (Sigma Aldrich, Darmstadt, Germany). Mouse embryonic stem cells (ESC) (LF2) were isolated from the mouse strain 129J [35] and originate from A. Smith (Oxford, UK). They were cultivated under feeder-free conditions on 0,1% gelantine-coating using knock-out Dulbecco’s Modified Eagle Medium (KO-DMEM) (Gibco, Carlsbad, CA, USA) supplemented with knock-out serum replacement (15%) (Gibco, Carlsbad, CA, USA), penicillin/streptomycin (1%), glutamax (1%), β-mercaptoethanol (5 × 10−5 M) (Invitrogen, Carlsbad, CA, USA) and leukemia inhibitory factor (LIF) (Millipore, Billerica, MA, USA) (1000 U/ml) at 37 °C in an atmosphere containing 5% CO2. b4-hiPSC were generated from human foreskin fibroblasts as described [36]. Cells were cultured on plates coated with reduced growth factor basement membrane matrix (Gibco, New York, NY, USA) in StemMacs medium (Miltenyi Biotec, Bergisch Gladbach, Germany), supplemented with 10 mM Y-27632 dihydrochloride (Sigma-Aldrich, St. Louis, MO, USA)

### Determination of cell viability

Cell viability was determined using the Alamar blue assay [37]. Viable cells are characterized by an effective mitochondrial metabolization of the non-fluorescent dye resazurin (Sigma, Steinheim, Germany) to fluorescent resorufin (excitation: 535 nm, emission: 590 nm). Relative viability in the untreated control was set to 100%. If not stated otherwise, data are shown as the mean ± standard deviation (SD) of ≥ three independent experiments each performed in biological quadruplicates. The combination index (CI) was determined [38] for the calculation of additive (CI >0.8<1.2), synergistic (CI≤0.8) or antagonistic (CI≥1.2) drug interactions in the co-treatment experiments.

### Analysis of doxorubicin import and export

Doxo import and export were measured by exploiting the inherent red fluorescence of Doxo as described [39]. Briefly, following 2 h of pulse-treatment with different Doxo concentrations, the fluorescence of the cells was measured as a surrogate marker of drug uptake by flow cytometry (excitation: 450 nm, emission: 560 nm). After a post-incubation period of up to 6 hours in the absence of Doxo, the residual intracellular fluorescence was again measured by flow cytometry. The time dependent decrease in Doxo fluorescence was calculated as a surrogate marker of Doxo export.

### Cell cycle analysis by flow cytometry

For flow cytometry-based analysis of cell cycle distribution, cells were trypsinized and combined with floating cells present in the medium. Cells were pelleted (1000 x *g*, 10 min, 4 °C) and suspended in PBS. DNase-free RNase A (Serva, Heidelberg, Germany) was added (2 μg/ml, 1h at RT). DNA was stained with propidium iodide (PI) (Sigma, Steinheim, Germany) for 20 min in the dark. Cell number was adjusted to 10^6^ cells/ml with PBS and analysis was performed using BD Accuri™C6 flow cytometer (BD, Franklin Lakes, NJ, USA).

### Analysis of apopotosis and senescence

Cell death by apoptosis was monitored by flow cytometry-based quantitation of the subG1 fraction, which represents the apoptotic cell fraction. Moreover, cleavage of PARP protein and pro-caspase-7 was monitored by Western blot analysis. To monitor senescence β-GAL expression was analyzed by Western blot analysis.

### Analysis of proliferation

To monitor proliferation, the incorporation of the nucleoside analogue 5-ethynyl-2′-deoxyuridine (EdU) into S-phase cells as well as the percentage of pH3 (Ser10) positive cells (mitotic index) and Ki-67 positive cells were determined. To this end, cells were seeded on cover slips and cultivated for the indicated time period. EdU-incorporation was analyzed using the EdU-Click 488 Kit (baseclick, Tutzing, Germany), which is based on a pulse-labeling of S-phase cells with 10 μM EdU according to the manufacturer’s protocol. To determine the mitotic index, cells were fixed with 4 % formaldehyde/PBS followed by incubation PBS-0.3 % TritonX-100 (5 min, RT). After blockage of unspecific binding (5 % BSA in 0.3 % Triton X-100/PBS (1 h, RT)), anti-Ser10 phosphorylated histone H3 antibody (pH3, Thermo Fisher Scientific Inc., Carlsbad, USA) (dilution 1:1000, 16 h, 4 °C) and Ki-67 antibody (Cell Signaling Technology Inc., Danvers, USA) (dilution 1:500, 16 h, 4°C) were added. Incubation with Alexa Fluor® 488 labeled goat anti-rabbit and Alexa Fluor 555 labeled goat anti-mouse secondary antibody was performed for 120 min at RT. pH3– and Ki-67-positive cells were counterstained with DAPI-containing Vectashield (Vector Laboratories, Burlingame, CA, USA) and analyzed by Olympus BX43 microscope (40 x objective) (Olympus, Hamburg, Germany).

### Immunocytochemistry-based analysis of DNA double-strand break (DSB) formation

The frequency of nuclear foci formed by S139 phosphorylated H2AX (γH2AX foci) is a commonly used surrogate marker of DNA double-strand breaks (DSBs) [40, 41] and was assayed by immunocytochemistry-based method. The appearance of nuclear 53BP1 foci, which is another marker of DSBs [42, 43], was also determined by immunocytochemistry. Upon treatment of cells grown on cover slips, cells were fixed with 4% formaldehyde in phosphate-buffered saline (PBS) (MERCK, Darmstadt, Germany) (15 min, RT) followed by incubation in PBS-0.3 % TritonX-100 (5 min, RT). After blocking (1 h, RT; blocking solution: 5% BSA (MERCK, Darmstadt, Germany) in PBS/0.3% Triton X-100 (Sigma, Steinheim, Germany)), incubation with γH2AX antibody (1:2000) and 53BP1 antibody (1:500) was performed overnight (4°C), followed by incubation with the secondary fluorescence-labelled antibody (1:500, 2 h, RT, in the dark). Cells were mounted in Vectashield (Vector Laboratories (Burlingame, CA, USA)) containing DAPI and the number of nuclear γH2AX foci and 53BP1 foci was scored (Olympus BX43 fluorescence microscope). Only nuclei with distinct foci were evaluated. γH2AX foci pan-stained nuclei, which are indicative of apoptotic cells, were excluded from the analyses. If not stated otherwise, data are shown as the mean ± SD from three independent experiments with each ≥50 nuclei analyzed per experimental condition.

### Western Blot Analysis

The activation of the DDR was investigated by Western blot analysis using total cell extracts obtained by lysing an equal number of cells in 150 µl RIPA buffer (20 min on ice). After sonication (EpiShear™ Probe sonicator, Active Motif, La Hulpe, Belgium) and centrifugation, Roti^®^-Load buffer (5 min, RT) was added to the supernatant and proteins were denatured by heating (5 min, 95°C). Afterwards, proteins were separated by SDS-PAGE (6 or 12.5% gel) and transferred onto a nitrocellulose membrane (Cytiva Europe GmbH, Freiburg im Breisgau, Germany) via the Protean Mini Cell System. After blocking (5% non-fat milk in TBS/0.1% Tween 20 (MERCK, Darmstadt, Germany) (2 h, RT)), the membrane was incubated with the corresponding primary antibody (1:1000, overnight, 4°C). After washing (TBS/0.1% Tween 20), the secondary (peroxidase-conjugated) antibody was added (1:2000, 2 h, RT). The ChemiDoc™Touch Imaging System (Bio-Rad Laboratories GmbH, Feldkirchen, Germany) was used for visualization of the bound antibodies.

### Quantitative mRNA expression analyses (RT-qPCR)

Total RNA was purified using the RNeasy Mini Kit (Qiagen, Hilden, Germany), followed by reverse transcriptase (RT) reaction with High Capacity cDNA Reverse Transcription Kit (Thermo Fisher Scientific Inc, Carlsbad, USA). For each PCR reaction 40 ng of cDNA and 0.25 µM of the corresponding primers (Eurofins MWG Synthesis GmbH, Ebersberg, Germany) were used. Quantitative real-time PCR analysis was performed in triplicates using the SensiMix SYBR Hi-ROX Kit (Meridian Bioscience, Cincinnati, USA) and a CFX96 Real-Time System (BioRad, Munich, Germany) with the Bio-Rad CFX Manager 3.1 software. PCR runs (35-40 cycles) were done as follows: 95°C – 10 min; 95°C –15 s; 60°C – 30 s; 72°C – 40 s; 72°C – 10 min. Melting curves were recorded to ensure the specificity of the amplification reaction. mRNA levels of *β-actin and GAPDH* were used for normalization. Unless stated otherwise, relative mRNA expression of untreated cells was set to 1.0.

### Statistical analyses

The Student’s t-test and the One-way ANOVA with Dunnett’s post-hoc test were employed to confirm statistically significant differences between different experimental groups. p≤0.05 was considered a statistically significant difference.

## Results and Discussion

### 1. Doxorubicin-resistant (A2780ADR) ovarian cancer cells are cross-sensitive to the anticancer drugs etoposide and cisplatin

We employed parental ovarian cancer cells (A2780) and thereof derived Doxo resistant variant (A2780ADR) for our study. Comparative analysis of their viability following Doxo treatment for 72 h revealed IC_50_ of 0.04 µM and 0.38 µM for A2780 and A2780ADR, respectively **(Fig. 1A).** A2780ADR cells revealed a profound cross-resistance to the Topo II inhibitor etoposide (Eto) (IC_50_ A2780: 0.14 µM; IC_50_ A2780ADR: 0.92 µM) **(Fig. 1B)**, while being only weakly cross-resistant to the platinating agent cisplatin (CisPt) ((IC_50_ A2780: 0.62 µM; IC_50_ A2780ADR: 1.91 µM) **(Fig. 1C)**. Based on the corresponding IC_50_ A2780ADR cells are characterized by a ∼10-fold, ∼7-fold and ∼3-fold higher resistance to Doxo, Eto and CisPt, respectively, as compared to the parental A2780 cells.

**Figure 1:**
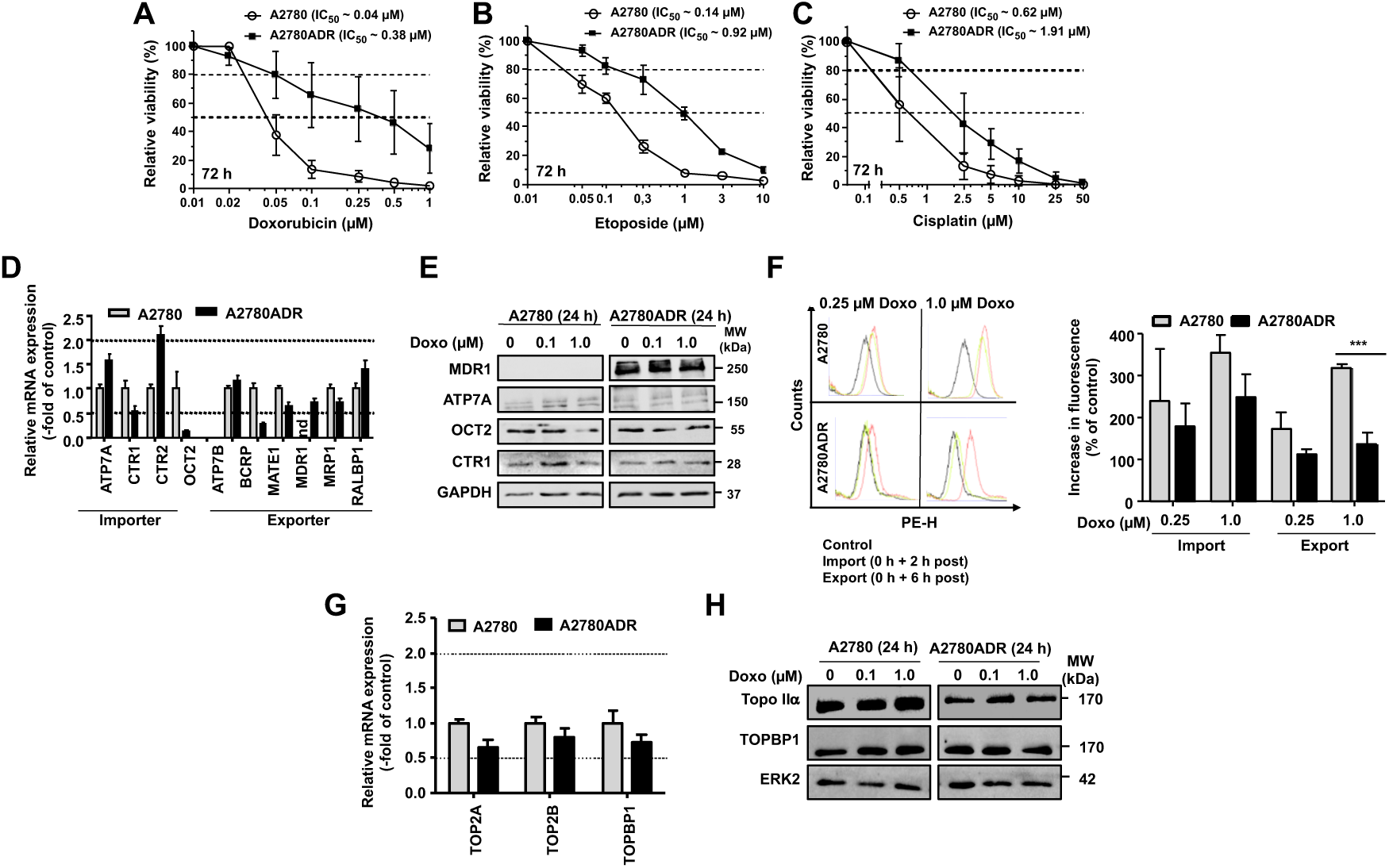
Comparative analysis of the response of A2780 and A2780ADR cells to treatment with anticancer drugs (Doxo, Eto and CisPt) and impact of mechanisms of drug transport. **A-C:** Logarithmically growing parental A2780 and A2780ADR variant cells were treated with the anticancer drugs doxorubicin (Doxo) (A), etoposide (Eto) (B) and cisplatin (CisPt) (C) at the indicated concentrations. 72 h after drug addition viability was monitored by use of the AlamarBlue assay as described in methods. Data shown are the mean ± SD from three independent experiments each performed in biological quadruplicates (n=3; N=4). Dashed lines indicate inhibitory concentrations (i.e. IC_20_ and IC_50_). For viability data (IC_50_) after 24 h and 72 h of treatment also see **Supplementary Fig. 1 and Supplementary Table 1**. **D:** Comparative analysis of the mRNA expression of selected transporters in A2780 and A2780ADR. Data shown are the mean ± SD from triplicate determinations. mRNA expression of transporters was normalized to GAPDH mRNA leves and set to 1.0 in the parental A2780 cells. The dashed lines indicate changes in mRNA levels of ≥ 2.0 and ≤ 0.5, which are considered as biologically relevant. Nd, not detectable. **E:** Comparative analysis of the protein expression of representative drug transporters under basal situation and after 24 h treatment with Doxo (0.1 µM, 1.0 µM). Data shown are from a representative western blot using ERK2 protein levels as loading control. Data obtained after 72 h are presented in **Supplementary Fig. 2**. **F:** Intracellular Doxo fluorescence was measured by flow cytometry-based methods after 2 h Doxo pulse-treatment (0.25 µM, 1.0 µM) and was taken as indicative of drug import. To measure drug export, Doxo pulse-treated cells were post-incubated for 6 h in the absence of the drug before fluorescence was monitored. Data shown in the left panel are representative results obtained form flow cytometry analyses. The histogram in the right panel depicts quantitative data obtained from n=3 independent experiments each performed in biological triplicates (N=3). Statistical significance: *** p≤0.001. **G:** Analysis of basal mRNA expression of topoisomerase II isoforms Topo IIα (TOP2A), Topo IIβ (TOP2B) and Topoisomerase binding protein (TopBP1). Data shown are the mean ± SD from triplicate determinations. Relative mRNA level in A2780 cells was set to 1.0. The dashed lines indicate changes in mRNA levels of ≥ 2.0 and ≤ 0.5, which are considered as biologically relevant. **H:** Comparative analysis of the protein expression of Topo IIα and TopBP1 under basal situation and after 24 h treatment with Doxo (0.1 µM, 1.0 µM). Data shown are from a representative western blot analysis using ERK2 as protein loading control. Data obtained after 72 h Dox treatment are presented in **Supplementary Fig. 2**.

### 2. Doxorubicin-resistant (A2780ADR) ovarian cancer cells are characterized by altered expression of drug transporters and enhanced doxorubicin export

Altered drug transport, as mediated for instance by p-glycoprotein, is one possible mechanism of acquired drug resistance of cancer cells [44, 45]. Analyzing the mRNA expression of various drug importers and exporters, we observed an elevated mRNA expression of the importer CTR2 and a reduced mRNA expression of the exporter BCRP in Doxo resistant A2780ADR cells as compared to the parental cells **(Fig. 1D).** Of note, while the mRNA expression of MDR1 was clearly detectable in A2780ADR cells, it was below detection limit in A2780 cells (**Fig. 1D**). Lack of MDR1 expression in the parental cells and high expression in A2780ADR was confirmed on the protein level **(Fig. 1E).** To monitor the cellś activity of drug import and export, we took advantage of the inherent redish fluorescence of Doxo and comparatively analyzed alterations in the intracellular fluorescence of A2780 and A2780ADR cells following Doxo treatment. The data show a similar increase in fluorescence in both cell lines after 2 h of Doxo pulse-treatment **(Fig. 1F)**, indicating that both cell lines have comparable Doxo uptake capacity. However, analyzing the remaining intracellular Doxo concentration after a subsequent 6 h post-incubation period in the absence of Doxo, residual fluorescence was significantly lower (∼50 %) in A2780ADR as compared to parental A2780 cells **(Fig 1F)**. This finding shows that A2780ADR cells are characterized by an about twice as fast drug export as A2780 parental cells, which is very likely due to their elevated MDR1 expression as concluded from the results of the mRNA and protein expression analyses. Thus, increased drug export likely contributes to the high Doxo resistance of A2780ADR cells. However, having in mind the ∼10-fold higher Doxo resistance of A2780ADR cells as compared to A2780 cells, we speculate that mechanisms others than just drug export additionally contribute to their pronounced drug resistant phenotype.

Since topoisomerase IIα and IIβ are relevant primary targets for the anticancer efficacy of Doxo, we additionally investigated their mRNA and protein expression. A2780 and A2780ADR cells revealed similar mRNA levels of Topo IIα, Topo IIβ and Topoisomerase IIβ binding protein (TopBP1) **(Fig. 1G).** However, on the protein level, A2780ADR were characterized by a reduced expression of Topo IIα protein and enhanced protein levels of TopBP1 under basal situation and 24 – 72 h after Doxo treatment **(Fig. 1H)**. Overall, this data indicates that both alterations in drug transport catalyzed by MDR1 and the protein expression of Topo IIα contribute to the strongly enhanced Doxo resistance of A2780ADR cells as well as to their profound cross-resistance to etoposide.

### 3. Analyses of proliferation and cell cycle progression of A2780 and A2780ADR cells following treatment with topoisomerase II inhibitors

To further analyze the netto outcome of altered drug export in A2780 versus A2780ADR cells, cell cycle progression was analyzed after 24 – 72 h of Doxo and Eto treatment. At early time point of analysis (i.e. 24 h), A2780 revealed a more pronounced accumulation of cells in G2/M phase as compared to A2780ADR **(Fig. 2A).** This effect was seen both upon Doxo and Eto treatment **(Fig. 2A)**. Of note, SubG1 fraction was not yet enhanced in A2780 cells at this time point of analysis. After an extended treatment period of 72 h, parental cells revealed a clear increase in the percentage of Doxo– and Eto-treated cells present in the SubG1 fraction, which was not observed in the drug resistant A2780ADR variants **(Fig. 2B).** Taken together this data shows that parental A2780 cells are hypersensitive to G2/M blocking activity and apoptosis induction triggered by Topo II inhibitory anticancer drugs.

**Figure 2:**
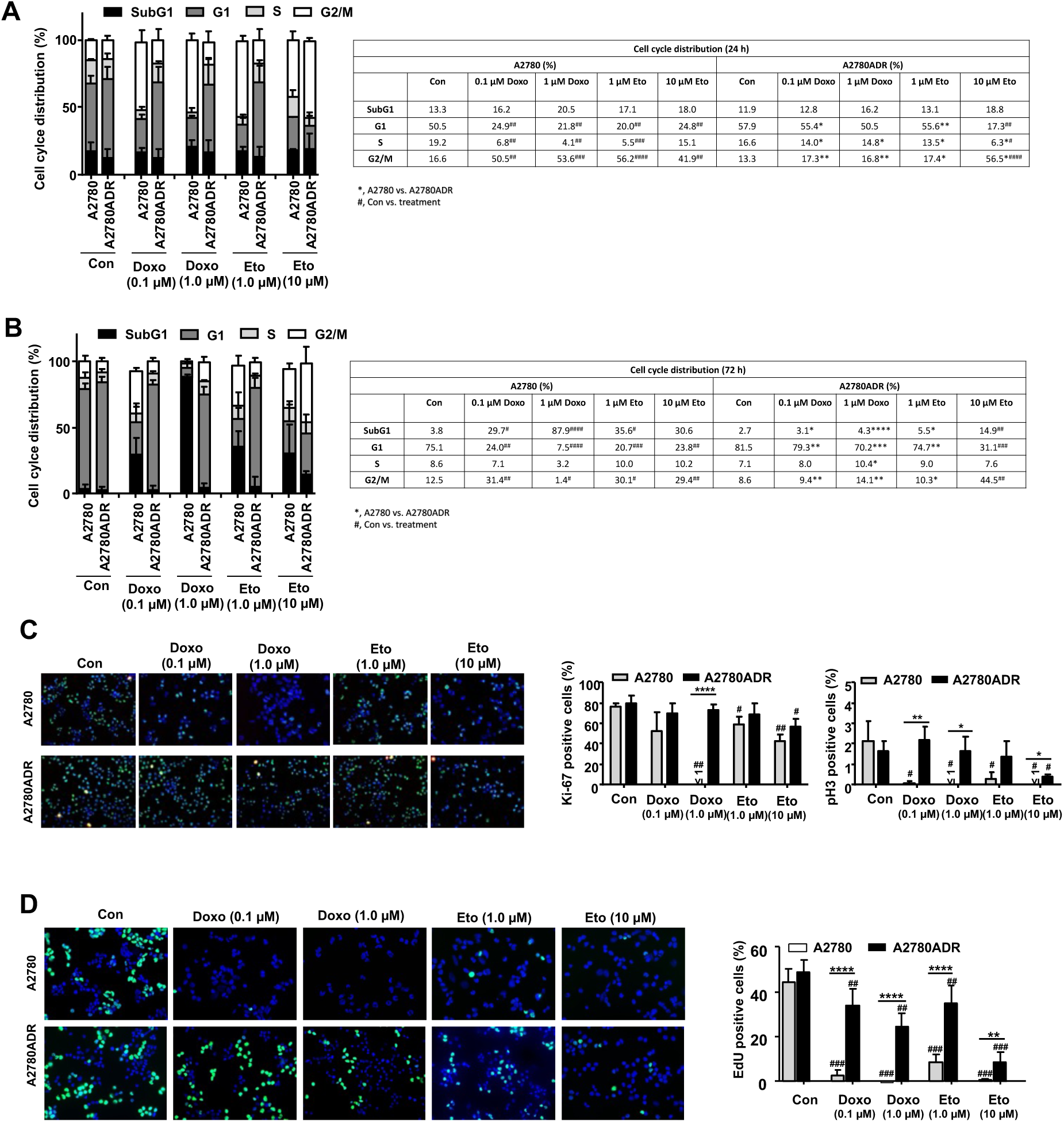
Cell cycle progression and proliferation following treatment of A2780 and A2780ADR cells with Topo II inhibitors. **A, B:** Logarithmically growing cells were treated with the indicated concentrations of doxorubicin (Doxo) or etoposide (Eto) for 24 h (A) or 72 h (B). Afterwards, cell cycle distribution was analyzed by flow cytometry and the percentage of cells present in different phases of the cell cycle (SubG1-, G1-, S– and G2/M-phase) was quantified. Data shown in the histogram (left panel) are the mean ± SD from n=3 independent experiments each performed in biological triplicates. The table on the right panel summarizes the mean values and indicates statistical differences between the individual groups. * p≤0.05; ** p≤0.01 (A2780 as compared to A2780ADR); ^#^ p≤0.05; ^##^ p≤0.01 (Con versus treated group) **C:** Logarithmically growing parental A2780 and Doxo resistant A2780ADR cells were treated with the indicated concentrations of Doxo (0.1 µM, 1.0 µM) or Eto (1.0 µM, 10 µM). 24 h later the percentage of Ki-67 positive or pH3 positive cells was determined as described in Methods. Quantitative data shown in the histogram are the mean ± SD from n=3 independent experiments, each performed with N=5 biological replicates. * p≤0.05; ** p≤0.01; **** p≤0.0001 (A2780 as compared to A2780ADR). Con vs. treatment: ^#^ p ≤ 0.05; ^##^ p ≤ 0.01; ^###^ p ≤ 0.001. **D:** Logarithmically growing parental A2780 and Doxo resistant A2780ADR cells were treated with the indicated concentrations of Doxo (0.1 µM, 1.0 µM) or Eto (1.0 µM, 10 µM). 24 h later cells were pulse-labeled with EdU for 2 h as described in Methods and the percentage of EdU positive cells was determined microscopically. Quantitative data shown in the histogram are the mean ± SD from five biological replicates. A2780 as compared to A2780ADR: ** p≤0.05; **** p≤0.0001. Con vs. treatment: ^##^ p ≤ 0.01; ^###^ p ≤ 0.001.

To monitor proliferative activity, the percentage of Ki-67 positive cells was analyzed in both cell variants. In addition, mitotic index was calculated by determining the percentage of pH3 positive cells. The data obtained show a profound decrease in the percentage of Ki-67 positive A2780 parental cells following treatment with Doxo but not Eto **(Fig. 2C).** By contrast, measuring the mitotic index, a significant decrease in the frequency of pH3 positive cells was found in both parental cells and resistant variants following Doxo or Eto treatment **(Fig. 2D)**. Measuring S-phase activity by analyzing the incorporation of EdU, we again observed a very strong reduction in the percentage of EdU positive parental cells upon both Doxo and Eto treatment, which was much weaker in the resistant A2780ADR cells **(Fig. 2D)**. Summarizing, the data show that A2780ADR cells are highly resistant to the antiproliferative activity of topo II inhibitory compounds.

### 4. Comparative analyses of Doxo-induced formation of DSB and activation of DDR-related mechanisms in A2780 and A2780ADR cells

Inhibition of topo II isoforms leading to the formation of DSB is considered as a major molecular mechanism underlying the anticancer activity of Doxo [46, 47]. Measuring the formation of nuclear γH2AX foci as well as 53BP1 foci and co-localized γH2AX/53BP1 foci, we observed a significantly lower number of DNA damage-associated foci in the A2780ADR cells as compared to the parental A2780 cells, both after treatment with Doxo and Eto **(Fig. 3A)**. Following high-dose treatment with Doxo, a high number of γH2AX pan-stained A2780 cells was detectable, which was not observed in A2780ADR cells **(Fig. 3A)**. The data show that the level of Doxo– and Eto-induced DSB is substantially reduced in the drug resistant variant.

**Figure 3:**
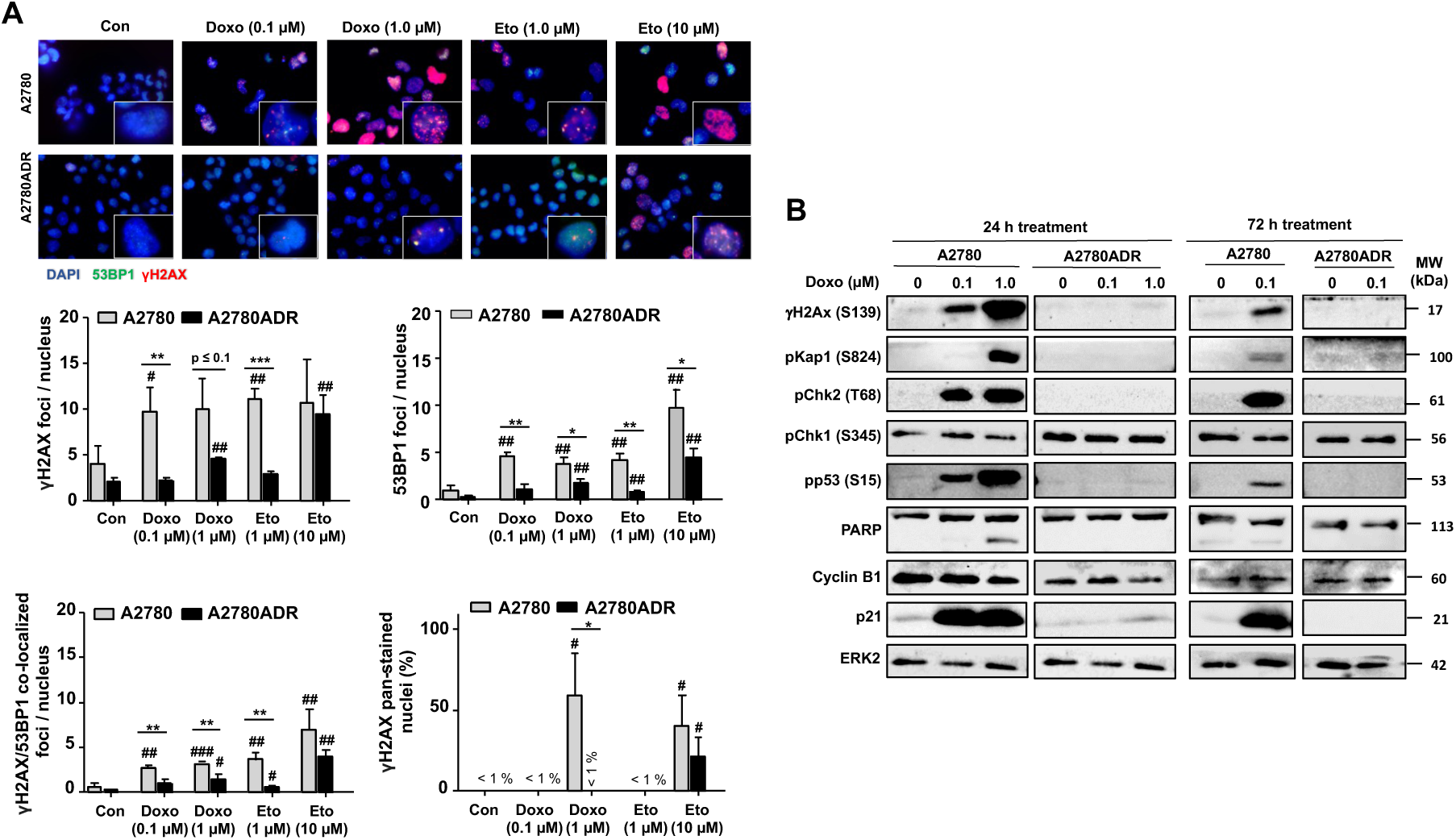
Impact of Topo II inhibitors on DNA damage formation and activation of DDR-related mechanisms in A2780 and A2780ADR cells. **A:** 24 h after treatment of logarithmically growing cells with the indicated concentrations of Doxo or Eto, the number of nuclear γH2AX-foci, 53BP1-foci, γH2AX/53BP1 co-localized foci and γH2AX pan-stained cells was analyzed. The upper part of the figure shows representative pictures. Quantitative data depicted in the histogram are the mean ± SD from n=3 independent experiments with each five pictures being analyzed per experimental condition. * p ≤ 0.05; ** p ≤ 0.01; *** p ≤ 0.001 (A2780 vs. A2780ADR); ^#^ p ≤ 0.05; ^##^ p ≤ 0.01; ^###^ p ≤ 0.001 (treated vs. untreated control). **B:** Logarithmically growing cells were treated with the indicated concentrations of Doxo for 24 h or 72 h. Afterwards, the protein expression of DDR-related factors was analyzed by Western blot analysis using EKR2 protein expression as loading control.

Having in mind that DSB are a potent trigger of mechnisms of the DDR, we next investigated the activation status of prototypical markers of the DDR by western blot analysis. Here, we observed an excessive increase in the DDR-related protein levels of γH2AX, pKap1, pChk2 and pp53 in Doxo treated A2780 parental cells only, both after Doxo treatment period of 24 h and 72 h. By contrast activation of these DDR-related factors was not found in A2780ADR cells **(Fig. 3B**). Of note, parental cells were also characterized by an enormous increase in the protein expression of the senescene marker p21, which was not detectable in the Doxo resistant A2780ADR cells **(Fig. 3B)**. Overall, these results support the hypothesis that the immense Doxo resistance of A2780ADR cells (i.e. ∼10-fold as compared to A2780) is also attributable to an attenuated Doxo-induced formation of DNA damage and largely reduced activation DDR-related mechanisms [48–50].

### 5. Cross-sensitivity of parental (A2780) and doxorubicin-resistant (A2780ADR) ovarian carcinoma cells to various anticancer drugs and inhibitors of DNA repair and DNA damage response (DDR)

Aiming to overcome the Doxo resistance of A2780ADR cells, we investigated their response to selected inhibitors of DDR– and DNA repair-related mechanisms, which are considered as promising targets to improve anticancer therapy and to overcome acquired drug resistance of tumor cells [49–52]. To this end, we comparatively investigated the outcome of a 72 h treatment period of A2780 parental and A2780ADR cells with Poly (ADP-ribose) polymerase (PARP) inhibitors (olaparib, niraparib), checkpoint kinase 1/2 (Chk1/2) inhibitors (prexasertib, rabusertib) and histone deacetylase (HDAC) inhibitors (entinostat, ricolinostat). In addition, the calcium channel blocker verapamil, which is reported to interfere with drug transport [53], the catalytic topo II inhibitor dexrazoxane, which is able to protect the heart from Doxo-induced cardiotoxicity by depleting both topo II isoforms independent of metal chelation [54–56] and Rac1 GTPase inhibitors (EHT1864, Ehop16), which are reported to interfere with Doxo-induced DNA damage formation and DDR activation [57, 58], were included into the study. As concluded from the IC_50_ determined, the data revealed a clear cross-resistance of A2780ADR cells to the PARPi olaparib (>10-fold) and niraparib (>5-fold), the Chk1/2i prexasertib (∼4-fold) and rabusertib (∼2.5-fold) and the HDAC class I inhibitor entinostat (∼3-fold) **(Fig. 4)**. A2780ADR cells were neither appreciably cross-sensitive nor hypersensitive to any of the other pharmacological modulators (i.e. ricolinostat, verapamil, EHT1863, EHOP16, dexrazoxane) employed. Hence, we conclude that targeting of single DNA repair/DDR-related mechanisms in anticancer drug-resistant A2780ADR cells by mono-treatment is not able to overcome their anticancer drug resistant phenotype. Moreover, based on the data obtained from the cross-resistance study, we hypothesize that mechanisms of DNA repair / DDR contribute to the Doxo-resistant phenotype of A2780ADR cells.

**Figure 4:**
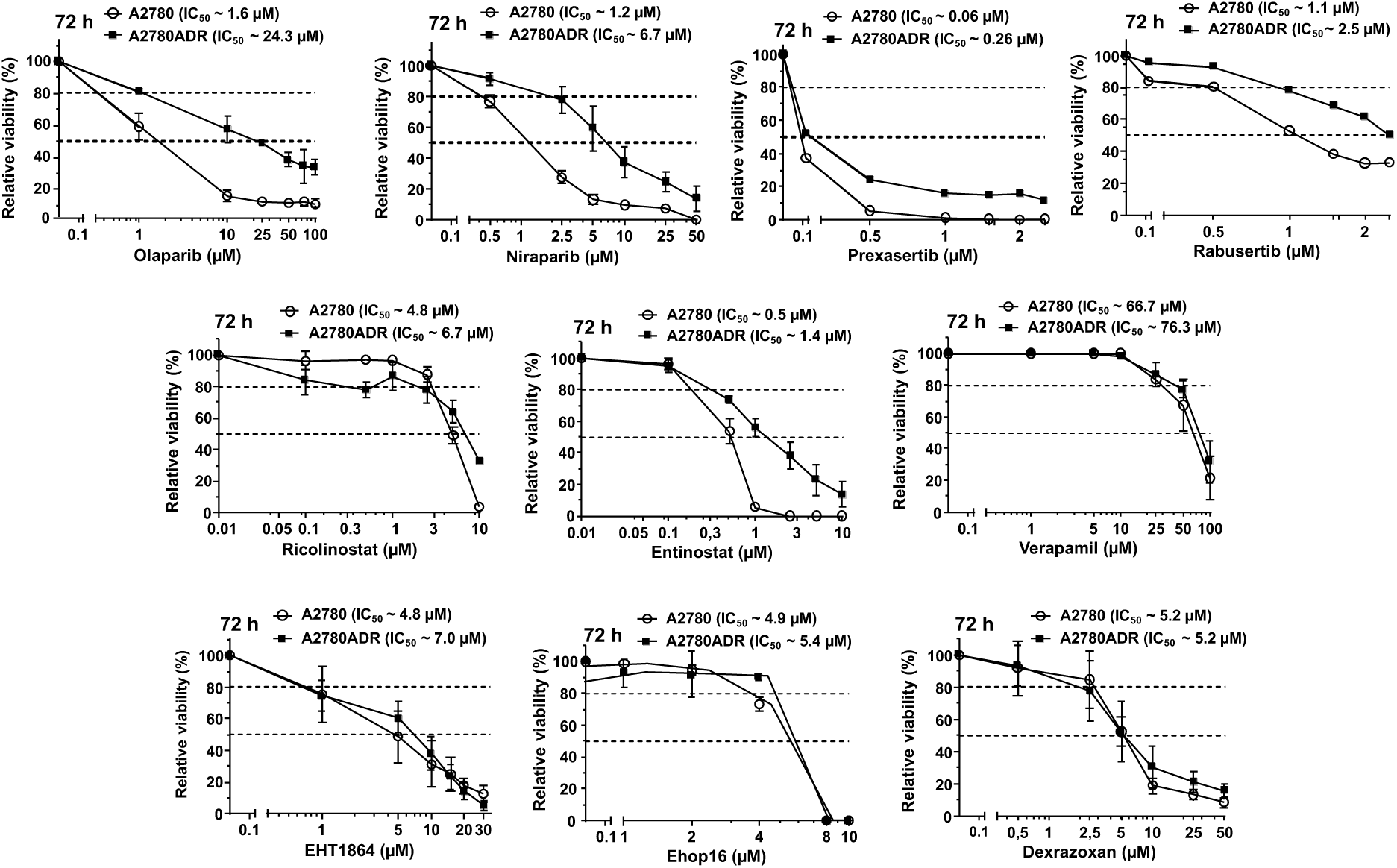
Analysis of cross-sensitivity of parental A2780 and Doxo-resistant A2780ADR cells to selected inhibitors of DDR– and DNA repair-related mechanisms. Logarithmically growing parental A2780 and A2780ADR variant cells were treated with selected pharmacological inhibitors of DNA repair (olaparib, niraparib), DDR (prexasertib, rabusertib), HDAC (ricolinistat, entinostat), Rac1 GTPase (EHT1864, Ehop16), drug transport (verapamil) and Topo II (dexrazoxane) at the indicated concentrations. 72 h after drug addition, viability was monitored by use of the AlamarBlue assay as described in methods. Data shown are the mean ± SD from three independent experiments each performed in biological quadruplicates (n=3; N=4). Dashed lines indicate inhibitory concentrations (i.e. IC_20_ and IC_50_). Data obtained from treatment period of 24 h are presented in **Supplementary Fig. 3.** For IC_50_ after 24 h and 72 h see **Supplementary Table 1.**

### 6. Co-treatment of doxorubicin resistant A2780ADR with DDR modifiers overcomes Doxo resistance by inducing synergistic toxicity

Based on the results of these extensive studies, we addressed the question whether the pharmacological inhibitors under investigation are able to overcome the acquired Doxo resistance of A2780ADR cells. To this end, entinostat, EHT1864, rabusertib, dexrazoxane and verapamil were selected for Doxo co-treatment analyses. Each of the selected inhibitors synergistically increased the cytotoxicity of Doxo in the drug resistant A2780ADR cells as concluded from the calculated combination indices (CI) **(Fig. 5A)**. As concluded from the combination index (CI), the most pronounced synergistic toxicity was observed when Doxo was combined with the transport inhibitor verapamil (Ver) or the Rac1 inhibitor EHT1864 **(Fig. 5A)**. The high efficacy of Ver correlates with increased intracellular steady state levels of Doxo under situation of co-treatment **(Fig. 5B)**. This was not found if EHT or EST were used for Doxo co-treatment **(Fig. 5B)**. The data indicate that synergistic toxicity resulting from combined treatment of A2780ADR cells with Doxo and Ver is at least partially due to Ver-mediated inhibition of transport mechanisms leading to lower concentrations of intracellular Doxo. By contrast, synergism observed especially upon use of EST and EHT1864 is suggested to be independent of drug transport. Of note, our data are in line with published data showing that Rac1 in involved in chemoresistance [59, 60] and class I HDACi are useful of overcome acquired anticancer drug resistance [61–64].

**Figure 5:**
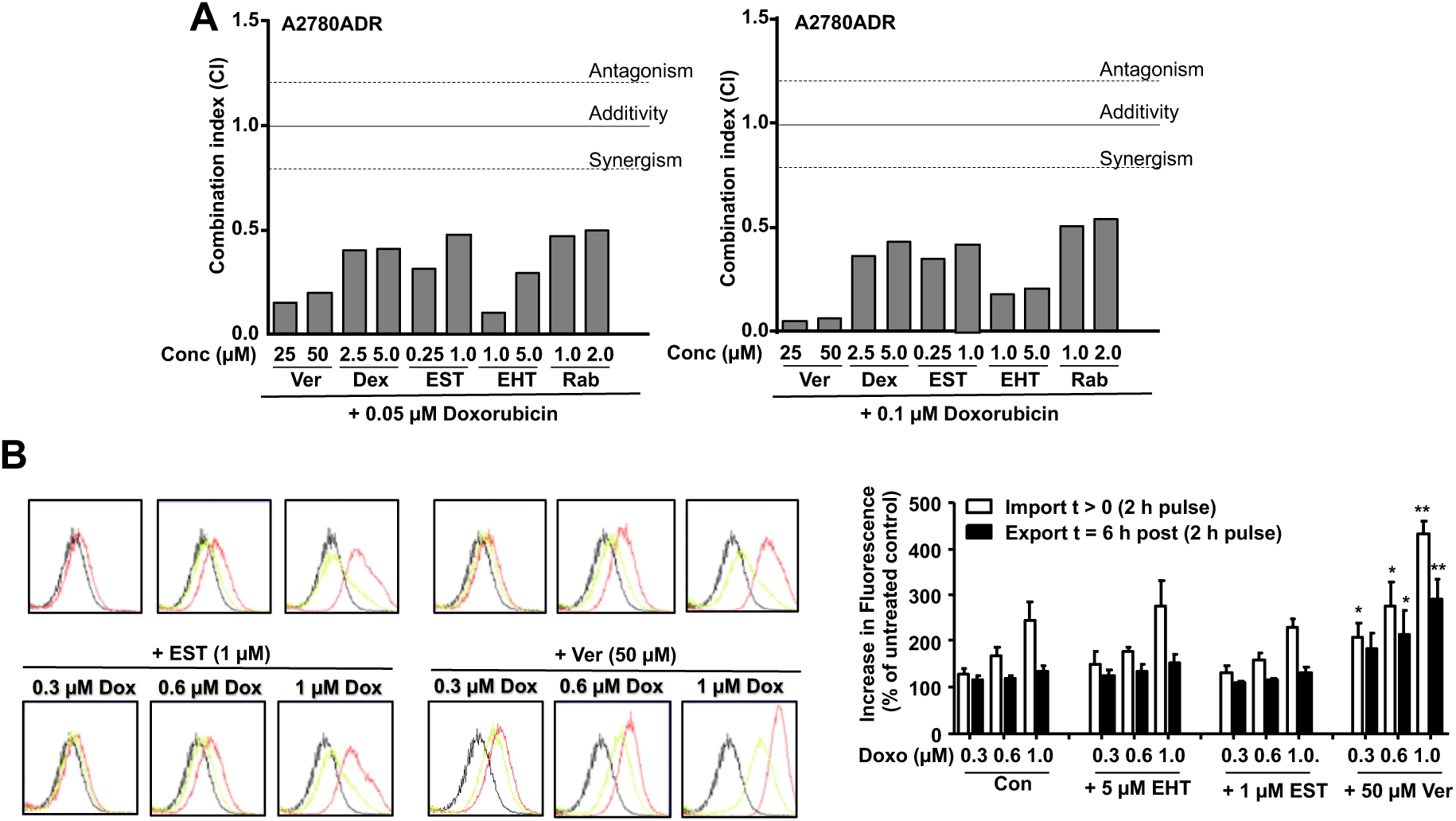
Combined treatment of A2780ADR with Doxo and selected inhibitors causes synergistic toxicity. **A:** Logarithmically growing doxorubicin resistant A2780ADR cells were co-treated with Doxo and selected pharmacological inhibitors at the indicated concentrations. 72 h after drug addition viability was monitored by use of the AlamarBlue assay and combination index (CI) was calculated as described in methods. Data shown are the mean ± SD from three independent experiments each performed in biological quadruplicates (n=3; N=4). **B:** Intracellular doxorubicin fluorescence was measured by flow cytometry-based methods after co-treatment with Doxo and selected pharmacological inhibitors as described in methods. To measure drug export, Doxo pulse-treated cells were post-incubated for 6 h in the absence of the drug before fluorescence was monitored. Data shown in the left panel are representative results obtained form flow cytometry analyses. The histogram in the right panel depicts quantitative data obtained from n=3 independent experiments each performed in biological triplicates (N=3). Statistical significance: * p≤0.05; ** p≤0.001.

Of note, Ver also synergistically increased the cytotoxicity of Doxo in A2780 parental cells **(Supplementary Fig. 4A**), showing the Ver effect is not restricted to cell with acquired Doxo resistance but also pertains to parental tumor cells. EST, EHT and Rab were also effective in combination with Doxo whereas Dex was not **(Supplementary Fig. 4A**). In addition, Ver also conferred synergistic toxicity in A2780ADR cells in combination with etoposide **(Supplementary Fig. 4B)**, showing that the Ver effect is not limited to Doxo but also comprises other anticancer drugs. Other inhibitors also conferred synergistic toxicity in combination with Eto, with EHT being most effective **(Supplementary Fig. 4B**). To further investigate whether Ver is also to overcome acquired resistance to anticancer drugs others than Topo II inhibitors, we investigated the influence of Ver on the CisPt sensitivity of CisPt-resistant A2780CisR cells, including EHT and EST for control. Data obtained show that all three modulators promoted CisPt-induced cytotoxicity in A2780CisR cells, with Ver and EST being most effective **(Supplementary Fig. 5)**. Based on this data we suggest that Ver is useful to re-sensitize both Doxo– and CisPt-resistant A2780 cells, indicating that Ver is able to overcome acquired drug resistance towards multiple anticancer agents.

In order to investigate whether the synergistic cytotoxicity evoked by combined treatment with Ver and Doxo is related to an elevated DNA damage induction, we measured the effect of the co-treatments on the formation of DSB. To this end, the number of nuclear γH2AX and 53BP1 foci, which are indicative of DSB, was analyzed. Data obtained show that Ver most significantly increased the number of both nuclear γH2AX and 53BP1 foci as well as γH2AX/53BP1 co-localized foci as compared to Doxo mono-treatment **(Fig. 6A)**. In addition, the percentage γH2AX pan-stained cells was also significantly enhance upon co-treatment with Ver and Doxo as compared to the mono-treatments **(Fig. 6A)**. Apparently, Ver is able to significantly increase the formation of DSB if used in combination with Doxo, demonstrating that Ver potentiates the genotoxic effects of Doxo in A2780ADR cells. Analyzing proliferation by monitoring the percentage of EdU positive cells, the most substantial antiproliferative effects were also observed upon combining Doxo with Ver **(Fig. 6B)**. In addition, Ver also significantly increased the percentage of PI positive cells if used in combination with Doxo **(Fig. 6C)**, showing that Ver potentiates Doxo-induced cell death. Summarizing this data, we suggest that Ver is able to overcome acquired Doxo resistance of A2780ADR cells by fostering the Doxo-induced formation of DSB leading to pronounced inhibition of proliferation and induction of cell death.

**Figure 6:**
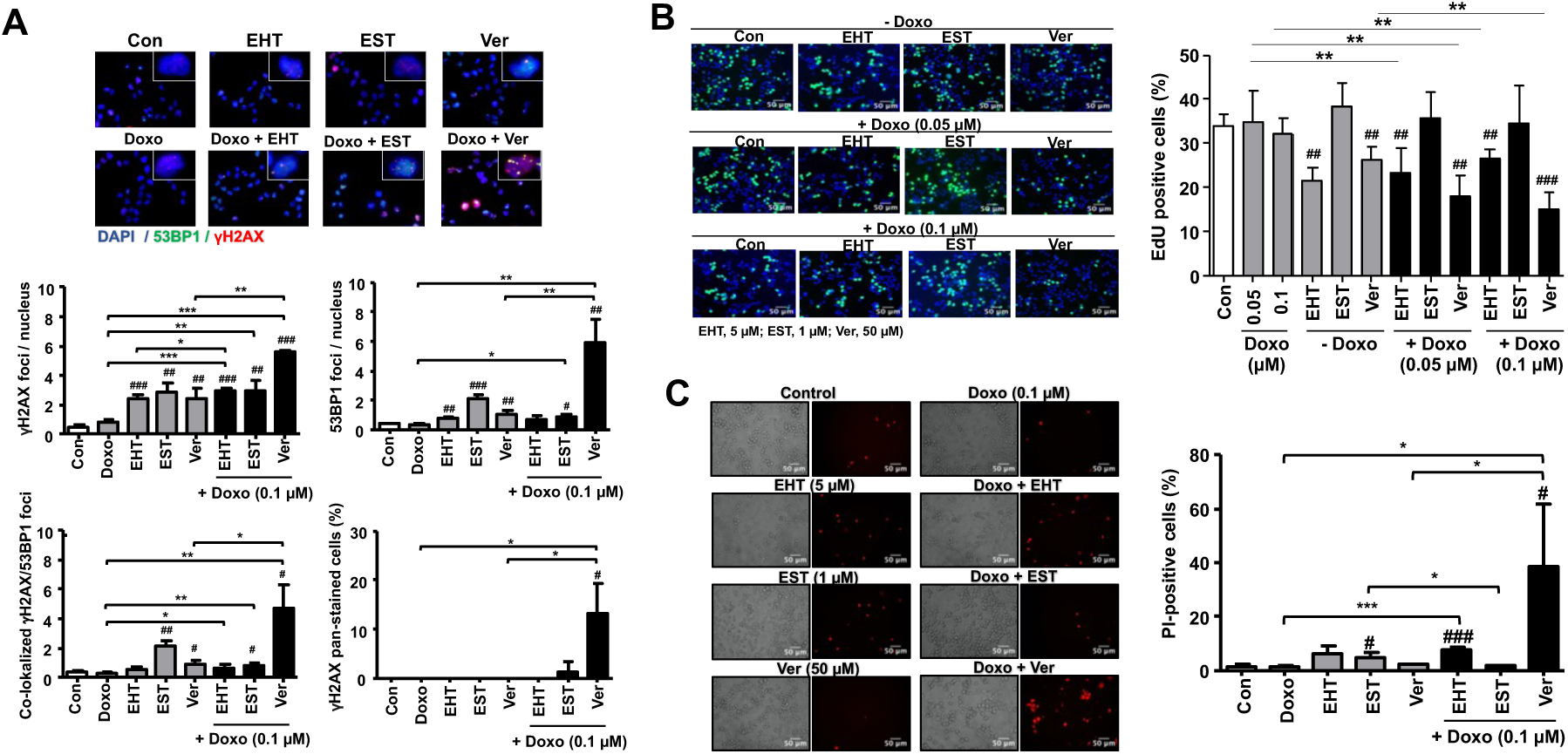
Influence of combined treatment of A2780ADR with Doxo and selected inhibitors on DNA damage formation, proliferation and cell death. **A:** 24 h after treatment of logarithmically growing cells with the indicated concentrations of Doxo and inhibitors (EHT, 5 µM; EST, 1 µM; Ver, 50 µM), the number of nuclear γH2AX-foci, 53BP1-foci, γH2AX/53BP1 co-localized foci and γH2AX pan-stained cells was analyzed as described in methods. The upper part of the figure shows representative pictures. Quantitative data depicted in the histogram are the mean ± SD from n=5 microscopical pics analyzed per experimental condition. Mono-treatment vs. co-treatment: * p ≤ 0,05, ** p ≤ 0,01, *** p ≤ 0,001); Untreated vs treated group: ^#^ p ≤ 0,05, ^##^ p ≤ 0,01, ^###^ p ≤ 0,001). **B:** Logarithmically growing Doxo resistant A2780ADR cells were co-treated with the indicated concentrations of Doxo and selected pharmacological inhibitors (EHT, 5 µM; EST, 1 µM; Ver, 50 µM). 24 h later, cells were pulse-labeled with EdU to monitor proliferation as described in Methods and the percentage of EdU positive cells was determined microscopically. Quantitative data shown in the histogram are the mean ± SD from five replicates. ** p≤0.05; **** p≤0.0001. ^##^ p≤0.01; ^###^ p≤0.001(as compared to untreated control). **C:** PI staining of mono– and co-treated A2780ADR cells 72 h after treatment with Doxo (0.1 µM) and pharmacological inhibitors (EHT, 5 µM; EST, 1 µM; Ver, 50 µM). Left panel: representative pictures; right panel: percentage of PI positive cells (mean ± SD from N=5 microscopical pics analyzed per experimental condition). Mono-treatment vs co-treatment: * p≤0.05; ** p≤0.01. Con vs treated group: ^#^ p≤0.05; ^##^ p≤0.01.

### 7. Influence of combined treatment of A2780 and A2780ADR cells with Doxo and selected inhibitors on mechanisms of the DDR and mRNA expression of selected susceptibility-related genes

Aiming to further elucidate the molecular mechanisms underlying the observed synergistic toxicity and to substantiate the increase in DSB formation under situation of Ver + Doxo co-treatment, we investigated the outcome of combined treatments on the activation status of a subset of DDR-related factors by western blot. The data obtained show that especially Ver is able to potentiate Doxo-mediated phosphorylation of both p53, RPA32, Chk2 and H2AX after 24 h or 72 h of co-treatment **(Fig. 7A)**. Moreover, if Doxo is combined with Ver, PAPR cleavage and cleavage of pro-caspase was observed **(Fig. 7A)**, pointing to the activation of apoptotic pathways under situation of co-treatment. Thus, we suggest that specifically inhibition of drug transport by Ver confers a broad re-sensitization of A2780ADR cells by fostering Doxo-induced formation of DSB and, in consequence, activation of DDR-related death associated mechanisms. Moreover, co-treatment of A2780ADR cells with Doxo + Ver increased the mean mRNA expression of selected factors involved in the regulation of oxidative stress response, DNA repair and cell cycle regulation as compared to Doxo or Ver mono-treated control **(Fig. 7B).** Taken together this data demonstrate that specifically Ver potentiates Doxo-stimulated activation of multiple DDR-related factors and alterations in the mRNA expression of susceptibility related factors.

**Figure 7:**
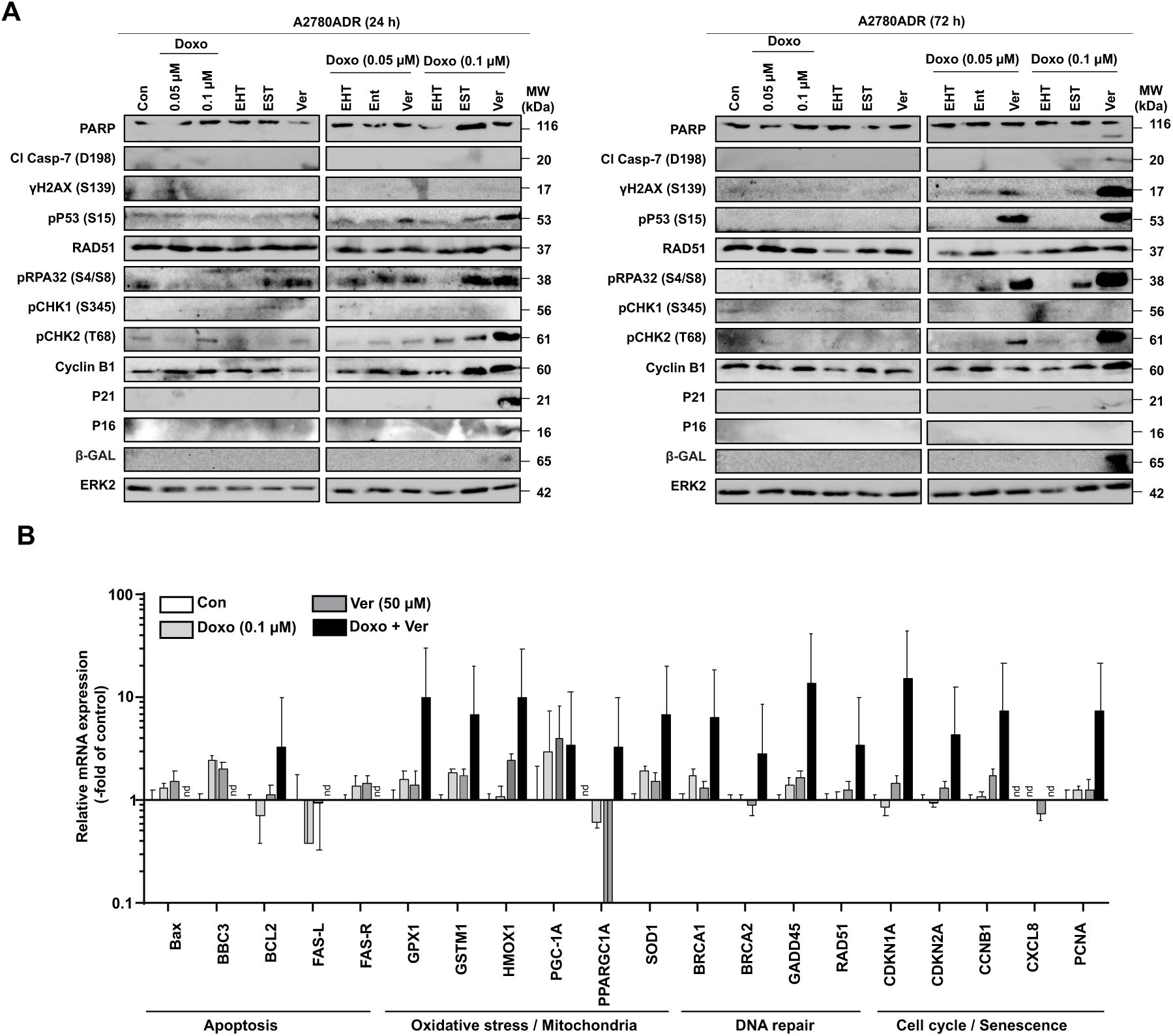
Influence of combined treatment of A2780ADR cells with Doxo and selected inhibitors on mechanism of the DDR and mRNA expression of selected susceptibility-related genes. **A:** Logarithmically growing A2780ADR cells were co-treated with the indicated concentrations of Doxo and selected pharmacological inhibitors (concentrations see Fig. 5) for 24 h or 72 h. Afterwards, the protein expression of DDR-related factors was analyzed by Western blot analysis. For loading control, blots were reprobed with ERK2 antibody. **B:** RT-qPCR-based analysis of the mRNA expression of selected factors known to contribute to different mechanisms of drug sensitivity. Data shown are mean ± SD from triplicate determinations as described in methods. Relative mRNA level in untreated A2780ADR cells was set to 1.0. nd, not detectable.

### 8. Influence of combined treatment on the viability of non-malignant cells

Irreversible cardiotoxicity is a dose-limiting adverse effect of anthracyclines [65, 66]. Hence, potentiating the anticancer efficacy of Doxo if combined with pharmacological modifiers brings up the concern of elevated side effects of the co-treatment, especially regarding heart damage. To address this aspect, we investigated the effect of combination treatments using immortalized murine HL-1 cardiomyocyte cells as an *in vitro* model. The results of this study revealed more than additive cytotoxicity if Doxo is combined with Ver **(Fig. 8A)**. By contrast, tendentially antagonistic effects were observed if Doxo was combined with the Rac1 inhibitor EHT1864 **(Fig. 8A)**. This is in line with previous studies showing that pharmacological inhibition of Rac1 is able to protect cardiac cell types from Doxo-induced injury *in vitro* and *in vivo* [67–70]. Moreover, we additionally investigated the cytotoxicity of Doxo in combination with Ver, EHT1864 and EST using stem cell lines of murine (mESC) and human (hiPSC) origin. We found that Ver promotes the Doxo sensitivity of both mESC and hiPSC **(Fig. 8B, 8C)**. Taken together, these data indicate that Doxo-induced toxicity might be elevated if used in combination with Ver, raising the concern of a possible increase in adverse effects resulting from co-treatment *in vivo*. Thus, while Doxo + Ver co-treatment is evoking substantial synergistic toxicity in Doxo resistant malignant cells, the therapeutic window of this combination treatment might be narrow. Noteworthy in this context, Rac1 inhibition by EHT1864 did not promote synergistic toxicity in combination with Doxo in non-malignant cells **(Fig. 8A-C)**. This finding is in line with *in vitro* and *in vivo* data [71–73] and points to a potentially favorable therapeutic window of EHT1864 if used in combination with Doxo.

**Fig. 8:**
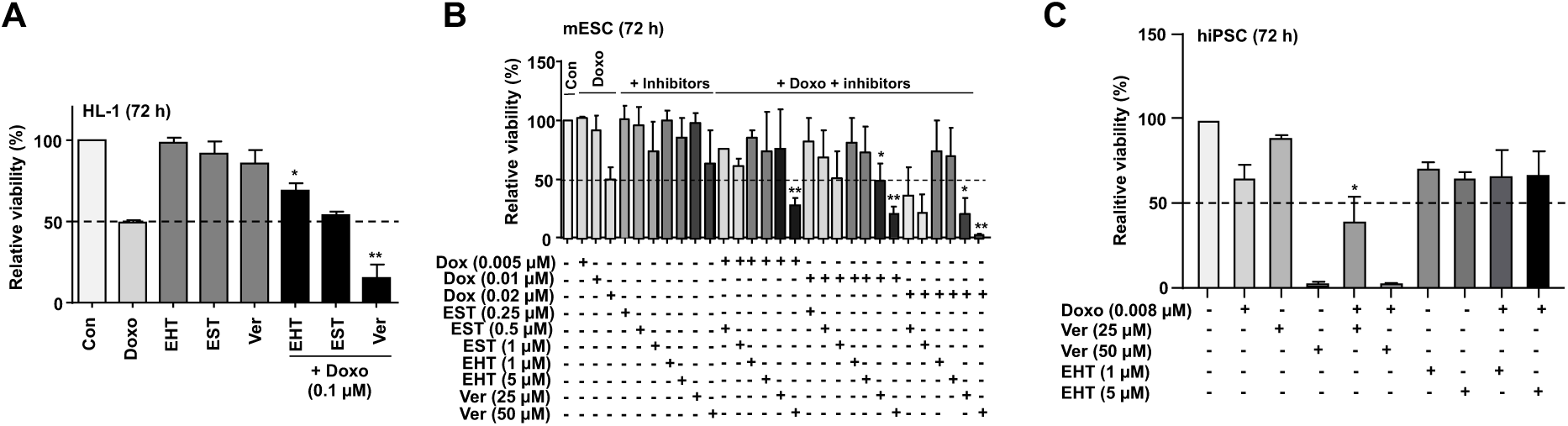
Effects of co-treatment of Doxo with selected pharmacological inhibitors on non-malignant cells. Logarithmically growing non-malignant murine HL-1 cardiomyocyte cells **(A)**, murine embryonic stem cells (mESC) **(B)** and human induced pluripotent stem cells (hiPSC) **(C)** were treated with the indicated concentrations of Doxo and selected pharmacological inhibitors for 72 h. Afterwards, cell viability was analyzed by the use of the AlamarBlue assay as described in methods. Data shown are the mean ± SD from n=1-3 independent experiments each performed in biological quadruplicates. * p≤0.05; ** p≤0.001 (mono-versus co-treated group). Dashed lines indicate 50 % viability.

Summarizing, the data indicate that acquired Doxo resistance of A2780ADR is associated with a reduced drug uptake due to high MDR1 expression and diminished Topo IIα expression. Furthermore, it is accompanied by an attenuated formation of DSB and reduced DDR activation as compared to A2780 parental cells. Acquired Doxo resistance of A2780ADR cells can be overcome most effectively by inhibition of drug export using verapamil (**Graphical abstract**). In consequence of Ver co-treatment, the intracellular steady state concentration of Doxo increases, promoting the formation of DSB, thereby triggering mechanisms of the DDR that inhibit cell proliferation and stimulate cell death. Noteworthy, targeting of Rac1– and/or HDAC-related mechanisms is also efficient to overcome acquired Doxo resistance independent of drug transport. Yet, the molecular mechanisms involved here are unclear and, hence, will be subject of forthcoming studies. In conclusion, we suggest that verapamil is highly effective to re-sensitize Doxo-resistant tumor cells, very likely by inhibiting MDR1-mediated drug efflux, thereby eventually potentiating the anticancer efficacy of Doxo. However, combined treatment with Ver may also amplify Doxo-stimulated adverse outcome pathways in non-malignant cells. Accordingly, forthcoming pre-clinical *in vivo* studies are needed to determine the therapeutic range of a combination treatment with Doxo plus verapamil.

## Abbreviations

ATM: Ataxia telangiectasia mutated
ATR: ATM– and Rad-3 related
BRCA 1/2: Breast cancer associated gene 1/2
cAT: conventional anticancer therapeutics
Chk1/2: checkpoint kinase 1/2 (CHEK1/2)
CisPt: cisplatin
DDR: DNA damage response
Doxo: doxorubicin
DSB: DNA double-strand breaks
EHT: Rac1 inhibitor EHT1864
Eto: etoposide
Est: entinostat
ERK2: extracellular regulated kinase 2
γH2AX: Ser139 phosphorylated histone H2AX
HDACi: histone deacetylase inhibitor
hiPSC: human induced pluripotent stem cells
HR: homologous recombination
Kap1: KRAB-associated protein 1 (TRIM28)
MDR1: multi-drug resistance gene 1
mESC: mouse embryonic stem cells
PARPi: Poly (ADP-ribose) polymerase inhibitor
Ver: verapamil

## Acknowledgments

This work was supported by the Deutsche Forschungsgemeinschaft (DFG Research Training Group (RTG) 417677437/GRK2578 (RG Fritz) and DFG Research Training Group (RTG) 270650915/GRK2158 (RG Fritz)).

## Author contributions

EM: Conceptualization, methodology, data generation, formal analysis, visualization, writing original draft

SF: Data generation, visualization

LM: Data generation, visualization

MS: Methodology, data generation, supervision

JM: Methodology, data generation

LA: Data generation, visualization

MUK: Resources, funding acquisition, writing original draft

GF: Conceptualization, funding acquisition, writing original draft, resources, supervision

