## Supplementary Figures and Table for "Overcoming acquired doxorubicin resistance of ovarian carcinoma cells by verapamil-mediated promotion of DNA damage-driven cell death"

Submitted to *Biorxiv*

### Supplementary data

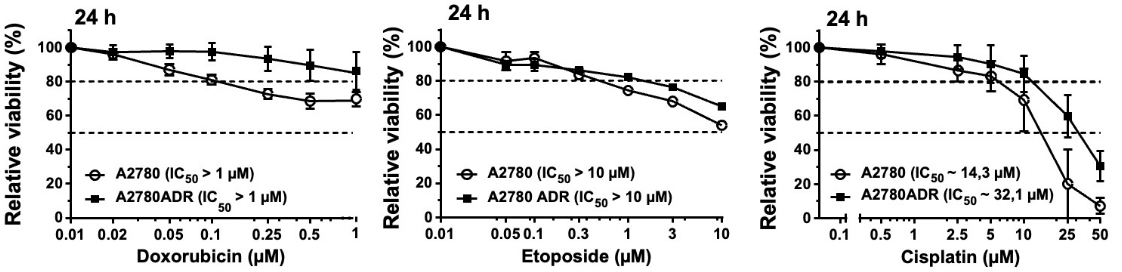


**Supplementary Fig. 1: Comparative analysis of the response of A2780 and A2780ADR cells to treatment with anticancer drugs (Doxo, Eto and CisPt).**

**A-C:** Logarithmically growing parental A2780 and A2780ADR variant cells were treated with the anticancer drugs doxorubicin (Doxo), etoposide (Eto) and cisplatin (CisPt) at the indicated concentrations. 24 h after drug addition viability was monitored by use of the AlamarBlue assay as described in methods. Data shown are the mean ± SD from three independent experiments each performed in biological quadruplicates (n=3; N=4). Dashed lines indicate inhibitory concentrations (i.e. $IC_{20}$ and $IC_{50}$).

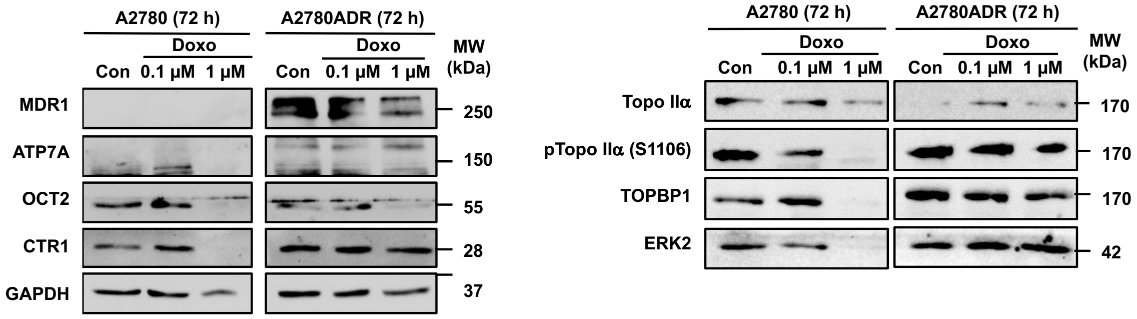


**Supplementary Fig. 2: Comparative analysis of the protein expression of selected transporters in A2780 and A2780ADR.**

Protein expression of selected drug transporters (left panel) and topoisomerase-related factors (right panel) was analyzed under basal situation and 72 h after treatment with Doxo (0.1 µM, 1.0 µM). Data shown are from a representative western blot. GAPDH and ERK2 protein expression were used as protein loading controls.

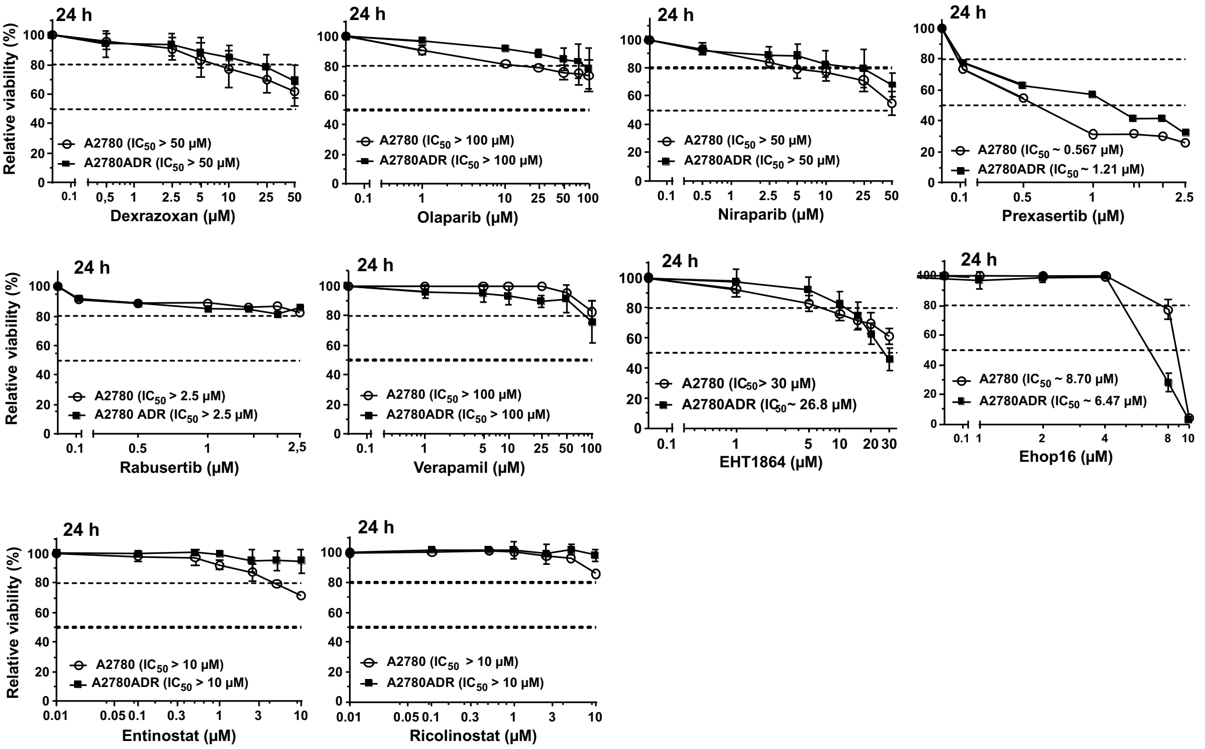


**Supplementary Fig. 3: Analysis of cross-sensitivity of A2780 and A2780ADR cells to selected pharmacological inhibitors of DDR- and DNA repair-related mechanisms.** Logarithmically growing parental A2780 and A2780ADR variant cells were treated with selected pharmacological inhibitors at the indicated concentrations. 24 h after drug addition viability was monitored as described in methods. Data shown are the mean ± SD from three independent experiments each performed in biological quadruplicates (n=3; N=4). Dashed lines indicate inhibitory concentrations (i.e. $IC_{20}$ and $IC_{50}$).

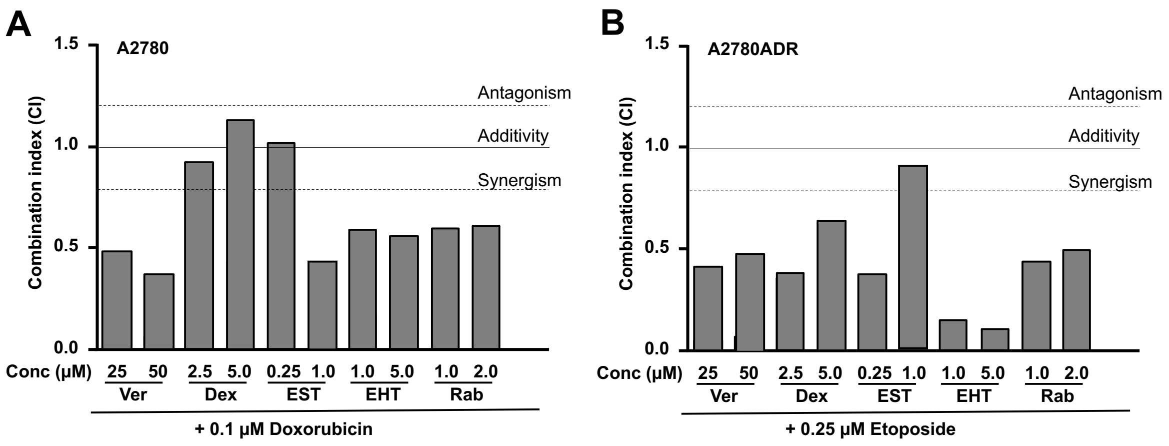


**Supplementary Fig. 4: Influence of pharmacological inhibitors on Doxo sensitivity of A2780 parental cells and Eto sensitivity of A2780 ADR cells.**

**A:** Logarithmically growing parental A2780 cells were co-treated with Doxo (0.1. µM) in combination with selected pharmacological inhibitors at the indicated concentrations. 72 h after drug addition viability was monitored by use of the AlamarBlue assay and combination index (CI) was calculated as described in methods. Data shown are the mean ± SD from three independent experiments each performed in biological quadruplicates (n=3; N=4).

**B:** Logarithmically growing doxorubicin resistant A2780ADR cells were co-treated with etoposide (Eto) (0.25 µM) in combination with selected pharmacological inhibitors at the indicated concentrations. 72 h after drug addition viability was monitored by use of the AlamarBlue assay and combination index (CI) was calculated as described in methods. Data shown are the mean ± SD from three independent experiments each performed in biological quadruplicates (n=3; N=4).

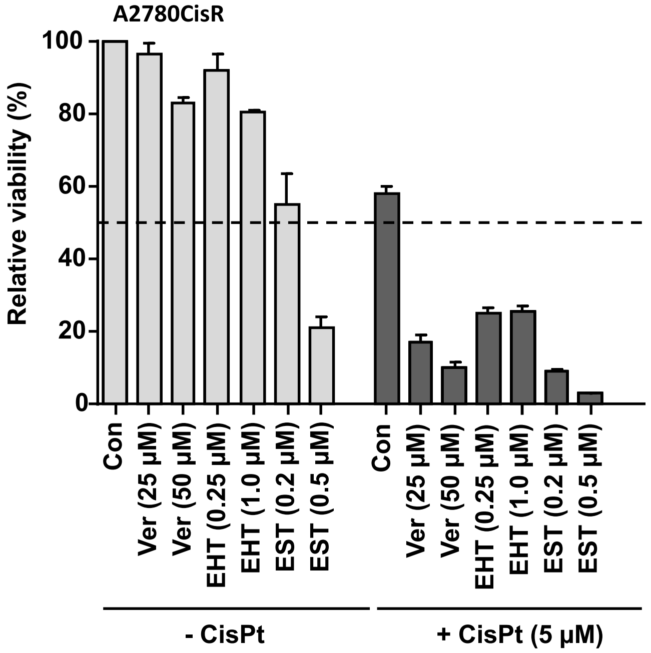


**Supplementary Fig. 5: Molecular effects of mono- and combined treatment of CisPt resistant ovarian A2780CisR cells with CisPt and selected inhibitors.**
Logarithmically growing Cisplatin (CisPt) resistant A2780CisR cells were left untreated (-CisPt) or were co-treated with CisPt (+CisPt) (5 µM) in combination with selected pharmacological inhibitors (Ver, EHT, EST) at the indicated concentrations. 72 h after drug treatment viability was monitored by use of the AlamarBlue® assay. Data shown are the mean ± SD from biological quadruplicate determinations. The dashed line indicates 50 % viability.

**Supplementary Table 1: Summary of the cross-sensitivity of A2780 and A2780ADR cells against selected anticancer drugs**

Cell viability of parental A2780 and Doxo resistant A2780ADR cells was analyzed after drug treatment period of 24 h or 72 h as described in methods (n=1-7 independent experiments; biological quadruplicates per concentration).

| Inhibitor | Experiments (N)* | Treatment period | Cell line | $IC_{50}$ (µM) | Resistance of A2780ADR |
|---|---|---|---|---|---|
| **Doxorubicin** | 7 | 24 h | A2780 | > 1 | 0 |
| | | 24 h | A2780ADR | > 1 | |
| **Etoposide** | 3 | 24 h | A2780 | > 10 | 0 |
| | | 24 h | A2780ADR | > 10 | |
| **Cisplatin** | 3 | 24 h | A2780 | ~ 11.5 | ++ |
| | | 24 h | A2780ADR | ~ 31.5 | |
| **Dexrazoxane** | 3 | 24 h | A2780 | > 50 | 0 |
| | | 24 h | A2780ADR | > 50 | |
| **Olaparib** | 3 | 24 h | A2780 | > 100 | 0 |
| | | 24 h | A2780ADR | > 100 | |
| **Niraparib** | 3 | 24 h | A2780 | > 50 | 0 |
| | | 24 h | A2780ADR | > 50 | |
| **Prexasertib (AZD77662)** | 1 | 24 h | A2780 | ~ 0.6 | + |
| | | 24 h | A2780ADR | ~ 1.2 | |
| **Rabusertib (LY2603618)** | 1 | 24 h | A2780 | > 2.5 | 0 |
| | | 24 h | A2780ADR | > 2.5 | |
| **EHT1864** | 5 | 24 h | A2780 | > 30 | 0 |
| | | 24 h | A2780ADR | ~ 26.8 | |
| **Ehop16** | 1 | 24 h | A2780 | ~ 8.7 | 0 |
| | | 24 h | A2780ADR | ~ 6.5 | |
| **Entinostat** | 2 | 24 h | A2780 | > 10 | 0 |
| | | 24 h | A2780ADR | > 10 | |
| **Ricolinostat** | 1 | 24 h | A2780 | > 10 | 0 |
| | | 24 h | A2780ADR | > 10 | |
| **Verapamil** | 2 | 24 h | A2780 | > 100 | 0 |
| | | 24 h | A2780ADR | > 100 | |

* Number of independent experiments
(each performed in biological quadruplicates)

| Inhibitor | Experiments (N)* | Treatment period | Cell line | $IC_{50}$ (µM) | Resistance of A2780ADR |
|---|---|---|---|---|---|
| **Doxorubicin** | 7 | 72 h | A2780 | ~ 0.04 | ++++ |
| | | 72 h | A2780ADR | ~ 0.40 | |
| **Etoposide** | 3 | 72 h | A2780 | ~ 0,14 | +++ |
| | | 72 h | A2780ADR | ~ 0.92 | |
| **Cisplatin** | 3 | 72 h | A2780 | ~ 0,4 | ++ |
| | | 72 h | A2780ADR | ~ 1,5 | |
| **Dexrazoxane** | 3 | 72 h | A2780 | 5.3 | 0 |
| | | 72 h | A2780ADR | 5.2 | |
| **Olaparib** | 3 | 72 h | A2780 | ~ 1.9 | ++++ |
| | | 72 h | A2780ADR | ~ 29.0 | |
| **Niraparib** | 3 | 72 h | A2780 | ~ 1.2 | +++ |
| | | 72 h | A2780ADR | ~ 6.8 | |
| **Prexasertib (AZD77662)** | 1 | 72 h | A2780 | ~ 0.06 | ++ |
| | | 72 h | A2780ADR | ~ 0.26 | |
| **Rabusertib (LY2603618)** | 1 | 72 h | A2780 | ~ 1.1 | + |
| | | 72 h | A2780ADR | ~ 2.5 | |
| **EHT1864** | 5 | 72 h | A2780 | ~ 4.8 | 0 |
| | | 72 h | A2780ADR | ~ 7.0 | |
| **Ehop16** | 1 | 72 h | A2780 | ~4.9 | 0 |
| | | 72 h | A2780ADR | ~ 5.4 | |
| **Entinostat** | 2 | 72 h | A2780 | ~ 0.5 | ++ |
| | | 72 h | A2780ADR | ~ 1,4 | |
| **Ricolinostat** | 1 | 72 h | A2780 | ~ 4.8 | 0 |
| | | 72 h | A2780ADR | ~ 6.7 | |
| **Verapamil** | 2 | 72 h | A2780 | ~ 66,7 | 0 |
| | | 72 h | A2780ADR | ~ 76,3 | |

0; considered as not resistant as compared to parental A2780 (difference $IC_{50}$ < 1.5)
+; weak drug resistance as compared to parental A2780 (difference $IC_{50}$ ≥ 1.5 < 2.5)
++; intermediate drug resistance as compared to parental A2780 (difference $IC_{50}$ ≥ 2.5 < 5.0)
+++; strong drug resistance as compared to parental A2780 (difference $IC_{50}$ ≥ 5.0 <10.0)
++++; very strong resistance as compared to parental A2780 (difference $IC_{50}$ ≥ 10.0)
